# Draft genome assembly of the green-bronze dung beetle, *Onthophagus orpheus*

**DOI:** 10.64898/2026.02.11.705363

**Authors:** Alexander J Bradshaw, Javier F. Tabima, Erin L McCullough

## Abstract

Dung beetles (Coleoptera: Scarabaeinae) are ecologically important insects, yet genomic resources for this diverse lineage remain limited. Here, we present a high-quality genome assembly for *Onthophagus orpheus*, an understudied species that is abundant in urban forests in the eastern United States. The assembled genome is highly contiguous and exhibits strong completeness, as assessed by Benchmarking Universal Single-Copy Ortholog (BUSCO) analyses, indicating robust representation of conserved protein-coding genes. Structural and functional annotation recovered a comprehensive gene set consistent with expectations for coleopteran genomes. This genome assembly provides an important resource for future work on the behavioral ecology and population genetics of *Onthophagus orpheus*, specifically, and Scarabaeinae more broadly.

## Introduction

Dung beetles (Coleoptera: Scarabaeidae: Scarabaeinae) are a diverse and ecologically important group of insects that provide essential ecosystem functions, including dung degradation, pest and parasite control, nutrient recycling, soil aeration, and secondary seed dispersal (Nichols et al. 2008). The dung beetle genus *Onthophagus* is particularly diverse, with over 2,000 described species worldwide (Emlen et al. 2005b). Members of this genus exhibit complex mating behaviors and intense male-male competition that have favored the evolution of sexually selected horns (Halffter and Edmonds 1982; Emlen and Philips 2006). Because of this morphological and behavioral variation, *Onthophagus* dung beetles have emerged as a model for studying the genetic and developmental mechanisms underlying phenotypic plasticity and evolutionary lability (Emlen et al. 2005a; Moczek 2005).

Despite their ecological and evolutionary significance, genomic resources for onthophagine dung beetles remain limited. Chromosome-level genome assemblies are available for *Onthophagus taurus, O. sagittarius, and Digitonthophagus gazella* (Davidson and Moczek 2024), but genomic resources are still lacking for other *Onthophagus* species. Generating additional high-quality genome assemblies will support population genetics studies and conservation efforts for this ecologically important lineage.

Here, we present a high-quality draft genome assembly for the green-bronze dung beetle *Onthophagus orpheus*. The species is native to the eastern United States and is the most abundant species in urban forests in central Massachusetts (McCullough et al. 2025). The assembly was generated using highly accurate PacBio HiFi sequencing. The resulting *O. orpheus* draft genomeimproves the genetic toolkit for this important genus and will facilitate comparative studies across Scarabaeinae.

## Results

### Sequencing Coverage, Genome Assembly, Contiguity, and Completeness

Genome sequencing depth was sufficient to support a high-quality assembly, with 99.45% of the assembly exhibiting normal coverage, indicating minimal under- or over-represented regions. Assembly accuracy was further supported by k-mer–based evaluation, which estimated 79.2% k-mer completeness and a consensus quality value (QV) of 59.9, corresponding to a very low base-level error rate. Structural correctness metrics were similarly high, with R-AQI and S-AQI scores of 93.9 and 97.7, respectively, indicating strong agreement between the assembly and underlying sequencing data.

The *O. orpheus* genome was assembled to a total length of 691.4 Mb, with the final assembly comprising 670 scaffolds, of which 667 exceed 10 kb, indicating a low degree of fragmentation. Assembly contiguity was high, with a scaffold N50 of 6.74 Mb and an N90 of 0.50 Mb, and the largest scaffold spanning 39.4 Mb. These metrics reflect strong long-range continuity and effective resolution of repetitive and structurally complex regions. Assembly completeness was also assessed using BUSCO with the coleoptera_odb12 dataset (n = 3,729). A total of 3,690 BUSCOs (98.9%) were recovered as complete, including 3,651 single-copy and 39 duplicated orthologs. Only 11 BUSCOs (0.3%) were fragmented, and 28 (0.8%) were missing. These results indicate that the assembly captures the vast majority of conserved beetle genes and is highly complete at the gene level. All total statistics are reported in Table 1.

**Table 1.** Genome assembly statistics and structural annotation for *Onthophagus orpheus*.

| Metric | Value |
| --- | --- |
| **Scaffolds** | **670** |
| **Scaffolds>10kb** | **667** |
| **Assembly size (MB)** | **691.4** |
| **Assembly N50 (MB)** | **6.74** |
| **Assembly N90 (MB)** | **50.3** |
| **Largest contig (bp)** | **39430979** |
| **Scaffolds >50 kb** | **518** |
| **% genome in scaffolds >50 kb** | **99.39** |
| **GC content (%)** | **33.8** |
| **Ambiguous bases (Ns)** | **0** |
| **BUSCO_db** | **coleoptera_odb12** |
| **BUSCO total genes** | **3729** |
| **BUSCO complete genes** | **3690** |
| **BUSCO complete single-copy genes** | **3215** |
| **BUSCO fragmented** | **11** |
| **BUSCO missing** | **28** |
| **merqury kmer completeness (%)** | **79.1993** |
| **merqury qv(phred)** | **59.9008** |
| **CRAQ R-AQI** | **93.86228757** |
| **CRAQ S-AQI** | **97.65696959** |
| **coverage normal(%)** | **99.4473** |
| **telomeric ends** | **0** |
| **telomeric ends(%)** | **0** |
| **Telomere to telomere scaffolds** | **0** |
| **Predicted genes** | **24840** |
| **Predicted transcripts** | **23317** |
| **Total exons** | **80476** |
| **Total repeat families** | **4022** |

### Gene Content, telomeric analysis, and Repetitive Elements

Structural annotation of the *O. orpheus* genome identified 24,840 predicted protein-coding genes, represented by 23,317 transcripts and 80,476 exonic regions. Analysis of telomeric features detected no telomeric repeat motifs using the ancestral insect motif “TTAGG” at scaffold termini, and no telomere-to-telomere scaffolds were recovered. Despite the absence of detectable telomeric ends, assembly contiguity was high, with 518 scaffolds exceeding 50 kb, accounting for 99.39% of the assembled genome.

Repetitive DNA analysis identified 4,022 repeat families across the genome. The assembled genome has a GC content of 33.8% and contains no ambiguous bases (Ns). Together, these results indicate a well-resolved gene set and repeat landscape suitable for downstream comparative and evolutionary analyses in Scarabaeinae.

### Functional annotation

Functional annotation of the *O. orpheus* genome was performed using a combination of similarity-based and domain-based approaches (see methods), resulting in putative functional assignments of predicted genes. In total, 25,294 protein-coding genes were predicted. Of these, 17,059 genes (67.4%) were assigned functional annotations based on eggNOG orthology, while 14,976 genes (59.2%) contained identifiable InterPro domains, and 12,679 genes (50.1%) were associated with Pfam protein families. Gene Ontology (GO) terms were assigned to 12,626 genes (49.9%), reflecting broad coverage of molecular functions, biological processes, and cellular components (Figure 2). Additionally, transmembrane and secreted proteins were predicted, and a large number were associated with membrane localization and transport: 4,090 proteins (16.2%) were predicted to contain transmembrane domains, and 2,439 proteins (9.6%) were predicted to be secreted. This highlights that O. orpheus has numerous gene families potentially involved in extracellular signaling, digestion, and environmental interaction.

**Figure 1.**
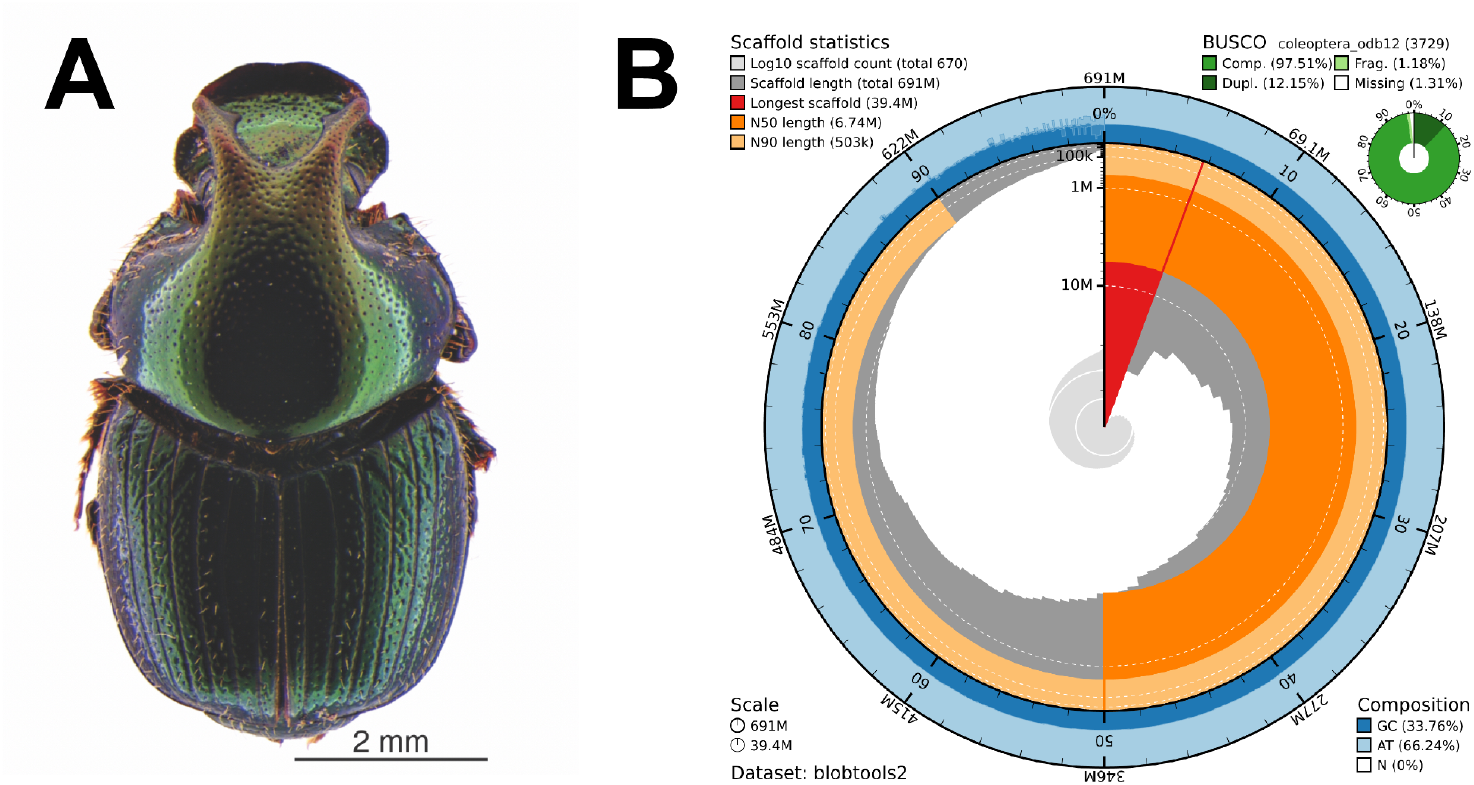
(A) Dorsal view of a large *Onthophagus orpheus* male Photo credit: Erin L McCullough. (B) Snail plot summarizing the genome assembly, including basid stats on base composition size as well as BUSCO results derived from the coleptera_odb12 database

**Figure 2.**
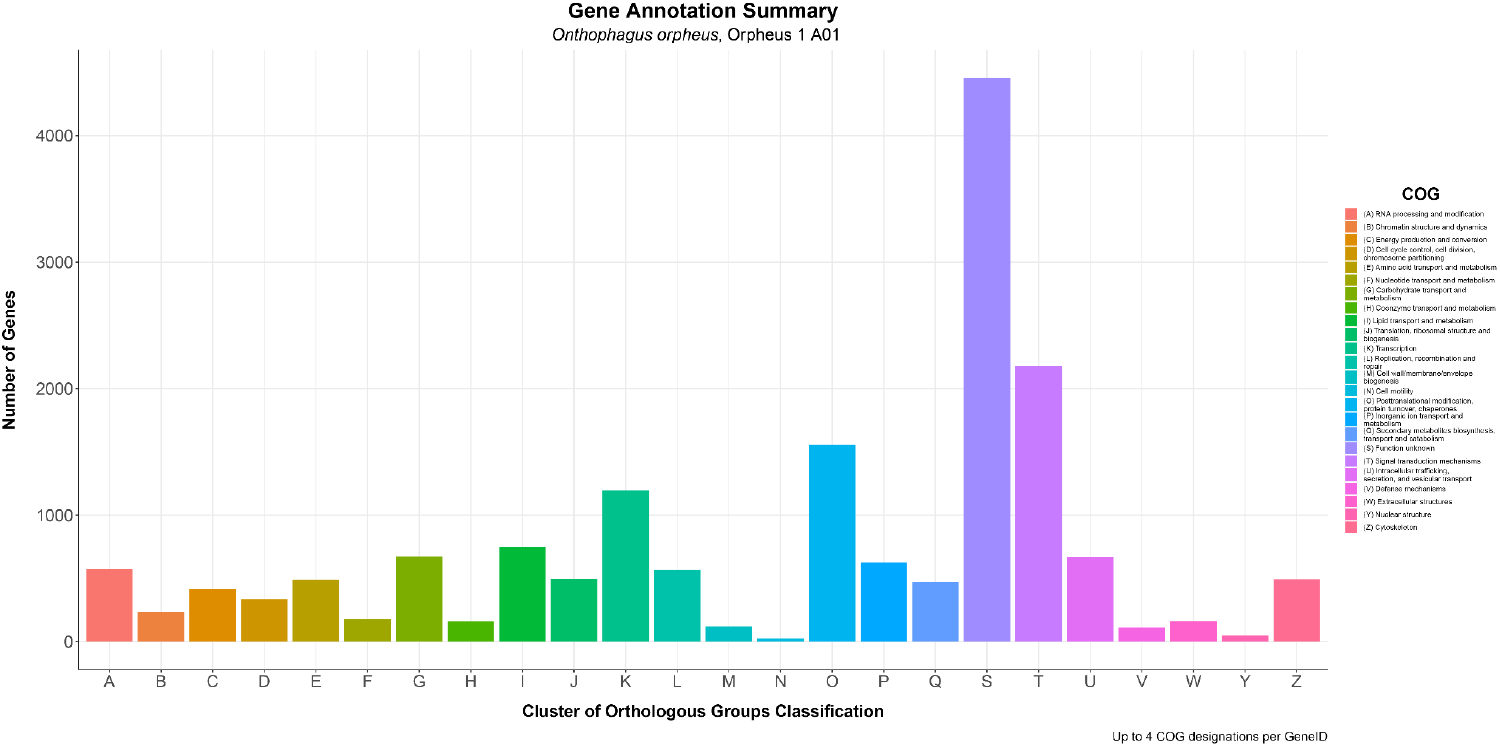
Gene ontology classification to Clusters of orthologous groups Clusters of orthologous groups are classified based on the Gene Ontology (GO) classification scheme. Each column and color along the x-axis represents a separate GO classification group, as described in the figure legend. The Y-axis represents the raw number of recovered genes assigned to each GO classification group.

Enzymatic gene families relevant to insect physiology and metabolism were also well recovered. Prediction and annotation identified 956 putative proteases and 301 carbohydrate-active enzymes (CAZymes), including glycoside hydrolases and carbohydrate esterases, which likely contribute to nutrient processing and digestion. Collectively, these results indicate that the O. orpheus genome annotation captures a diverse and functionally rich gene repertoire. All annotation data has been provided in the supplementary data.

## Discussion

This genome assembly provides a high-quality genomic resource for *Onthophagus orpheus*, an ecologically important dung beetle within the subfamily Scarabaeinae. The large, contiguous assembly and high BUSCO completeness indicate that the majority of protein-coding genes and conserved genomic regions are well represented, supporting the use of this genome as a reliable reference for downstream analyses. As such, the *O. orpheus* genome will be valuable for future comparative genomic studies within Scarabaeidae and Coleoptera, where sparse taxon sampling has constrained inference of genome evolution. Comparative analyses leveraging this genome can identify lineage-specific expansions, contractions, and structural variation associated with dung-feeding specialization, detoxification, and sensory adaptation, ecological processes known to drive diversification in beetles (McKenna et al. 2019).

The generation of a highly quality reference genome of *O. orpheus* also enables us to pursue future studies related to population-level analyses. High-quality reference genomes are essential for accurately mapping resequencing data, detecting selection, and reconstructing demographic history (Ellegren 2014), a growing research area within entomology (Raszick et al. 2021; Webster et al. 2023; Franzem et al. 2025). Dung beetles are sensitive to anthropogenic disturbance, including agricultural intensification, habitat fragmentation, and climate change, which can strongly influence population structure and adaptive variation (Davis et al. 2001; Nichols et al. 2007). This genome enables the identification of genomic regions associated with local adaptation, stress tolerance, and dispersal, and provides a framework for future conservation-genomic studies in dung beetles.

The *O. orpheus* genome assembly adds to a small but growing number of genomic resources for the speciose genus *Onthophagus*. This genus has emerged as a model system for studying sexual selection because of its remarkable diversity in horn morphology, condition-dependent trait expression, and alternative reproductive tactics. However, chromosome-level assemblies are currently available for only three species: *O. taurus, O. sagitarius*, and *Digitonthophagus* (formerly *Onthophagus*) *gazella* (Davidson and Moczek 2024). The availability of a reference genome for *O. orpheus* will facilitate comparative studies on the genomic architecture underlying horn development and horn polymorphisms (Kijimoto et al. 2013; Casasa et al. 2017), as well as investigations on mating systems and patterns of polyandry in natural populations (McCullough et al. 2017).

More broadly, this work improves the taxonomic representation of assembled beetle genomes and contributes to the study of insect genome evolution. Beetles exhibit extensive variation in genome size, repeat composition, and chromosomal organization, and comparative analyses suggest that transposable elements and gene duplication play major roles in shaping coleopteran genomes (Peters et al. 2017; McKenna et al. 2019). The *O. orpheus* assembly contributes to these efforts by providing a scarabaeine reference that can be integrated into large-scale comparative datasets to understand how ecological specialization and developmental novelty influence genome architecture.

Together, this genome assembly improves genomic representation within Scarabaeidae and provides a resource for future studies in understanding the biology of the genus *Onthophagus*. As additional dung beetle genomic data becomes available, the *Onthophagus orpheus* genome will serve as a key comparative reference for investigating the genomic basis of ecological specialization, developmental innovation, and the evolution of sexually selected traits in an ecologically impactful beetle lineage.

## Methods

### Specimen acquisition, genomic DNA extraction, and sequencing

Beetles were collected with pitfall traps on June 20, 2024, from Hadwen Arboretum (42.258°N, −71.832°W). We followed the basic salt-extraction method of Aljinabi and Martinez (1997) to extract genomic DNA from adult male *Onthophagus orpheus*. The legs, head, and thorax of each individual was homogenized in 350 mL sterile salt homogenizing buffer (50 mM Tris-HCl pH 8.0, 100 mM EDTA, 100 mM NaCl, and 1% SDS) and 2.5 mL of 20 mg/mL proteinase K using a pellet pestle. The samples were incubated at 65°C with intermittent vortexing for 15 minutes, cooled on ice, and mixed with 150 mL 5M NaCl. Samples were vortexed vigorously for 10s and spun down at 13000 rpm for 10 min. The supernatant was transferred to fresh tubes, mixed with 500 mL of isopropanol, and centrifuged at 13000 rpm for 10 min. The pellet was washed with 70% ethanol, dried, and finally resuspended in 50 ul TE buffer. Genomic DNA (13.56 ug) from one individual was sent to the UMass Chan Medical School Deep Sequencing Core for PacBio sequencing.

### Data quality control, Genome assembly, and quality assessment

Genome assembly of *Onthophagus orpheus* was performed using the PacBio HiFi Assembly pipeline developed by the Harvard Informatics Group (https://github.com/harvardinformatics/pacbio_hifi_assembly), which incorporates a series of optimized tools for processing and assembling PacBio HiFi reads using the highly accurate Hifiasm (Cheng et al. 2021) genome assembler into a reproducible Snakemake workflow (Köster and Rahmann 2012). The pipeline automates read correction, consensus generation, and assembly graph construction, providing a reproducible framework for high-quality long-read genome assemblies. All analyses were executed using a Singularity container cloned directly from the public repository and run with default parameters appropriate for diploid eukaryotic genomes to ensure software consistency and reproducibility. The resulting primary contig (Orpheus_1_A01.p_ctg.fa) served as the base assembly for subsequent quality control and contamination removal.

### Quality assessment, Contamination Removal, Pre-processing, and Repetitive region masking

Initial assembly quality and structural integrity were assessed using QUAST v5.2.0 (Gurevich et al. 2013), comparing the *O. orpheus* assembly to the *Onthophagus taurus* (GCF_036711975.1) and *Onthophagus sagittarius* (GCA_036711965.1) reference genomes, as well as the raw unpolished PacBio HiFi assembly (Supplementary data). Visualization of the whole genome and BUSCO scores was performed using the blobtk plot -v snail function of the blobtookit (Challis et al. 2020) suite of tools. Metrics, including contig count, N50, and total assembly length, were recorded to evaluate polishing and contamination-removal performance and are provided in the supplementary data.

Following assembly quality analysis, the genome was screened for potential adapter and foreign organismal contamination using the NCBI Foreign Contamination Screen (FCS-GX) toolset v0.5.5 (Astashyn et al. 2024). The goal was to identify and remove non-target sequences before downstream annotation. Screening was performed against the NCBI *Genome Taxonomy Database*, using the Onthophagus taxonomic identifier (NCBI:txid166331) as the reference for contamination removal.

The contamination-free assembly was then preprocessed for annotation using the “clean” function of Funannotate v1.8.17 (Palmer and Stajich 2020) to remove potential low-quality and redundant scaffolds, as well as those below 500 bp. Repetitive elements were identified using RepeatModeler v2.0.7 (Flynn et al. 2020), which performs de novo repeat discovery by integrating the RepeatScout (Price et al. 2005) and RECON (Bao and Eddy 2002) algorithms. This analysis identified 4,022 repeat families, which were subsequently used as a species-specific repeat library (Supplementary data) to mask repetitive regions of the assembly before gene prediction and annotation.

### Gene prediction and function annotation

For integrated gene prediction, the “predict” function of Funannotate v1.8.17 was employed using the masked repetitive-region assembly. Here, we used a combination of Augustus (Stanke et al. 2006), (Borodovsky and Lomsadze 2011), GlimmerHMM (Majoros et al. 2004), and SNAP (Korf 2004) gene prediction software with the flags --busco_seed_species tribolium and --protein_evidence fromthe protein sequences of the *O. taurus* reference (GCF_036711975.1) to guide gene prediction. High-quality predictions were then determined using EVidenceModeler (Haas et al. 2008), with equal weight given to all gene prediction methods and repeat regions incorporated into the EVM step (--repeats2evm) to improve gene model placement in masked regions. After prediction, the resulting Augustus model parameters were archived with the Funannotate “species” functionto preserve the optimized *O. orpheus* species profile for future gene prediction experiments, which are also provided in the supplementary data.

Functional annotation of predicted gene models was performed using the Funannotate “annotate: function, with the insecta_odb9 BUSCO database guiding quality evaluation. Functional Annotation was then performed with InterProScan5 5.59_91.0 (Jones et al. 2014), eggNOG-mapper v2.1.12 (Cantalapiedra et al. 2021). Finally, secreted and transmembrane protein prediction was performed with Phobium v1.01 (Käll et al. 2004) and the fast algorithm of SignalP 6 (Teufel et al. 2022).

### Quantification and visualization of the Final assembly and annotations

Quality assessment using the PAQman v1.2.0 (O’Donnell et al. 2025) pipeline, which leverages multiple commonly used and well-established bioinformatic tools to comprehensively assess genome assemblies. This includes: Contiguity is assessed using QUAST (Gurevich et al. 2013); BUSCO scores (Simão et al. 2015); completeness and base-level error rates estimation by Merqury (Rhie et al. 2020); Structural correctness metrics using Classification of Regional Assembly Quality (CRAQ) (Li et al. 2023); Coverage metrics generated through read mapping and utilities using Minimap2 (Li 2018) for long and short read alignment, Samtools for alignment processing (Danecek et al. 2021), and Bedtools (Quinlan and Hall 2010) for genomic coverage calculations. Finally, telomerality features are derived from sequence processing with Seqtk (Li 2026) and Bedtools (Quinlan and Hall 2010). Visualization of gene ontology results from the function annotation was performed by mutating the data frame output from Funannotate using the R package tidyverse v2.0.0 (Wickham et al. 2019), then plotting using ggplot2 v 4.0.0(Wickham 2016). Final assembly statistics are presented in Table 1, and all PAQman outputs are provided in supplementary data.

## Author Contributions

Alexander J Bradshaw performed data analysis, figure creation, and preparation of the manuscript. Javier F. Tabima performed data analysis and manuscript preparation; Erin L. McCullough performed specimen capture, DNA sequencing, figure creation, and manuscript preparation.

## Data availability

All Raw sequencing reads have been deposited in the NCBI Sequencing Read Archive (SRA) under Bioproject number PRJNA1395262. Raw sequencing reads are deposited on the Sequencing Read Archive (SRA) under the accession number SRR36630064. The Draft genome assembly and annotations are deposited under NCBI accession numbers xxxxx and xxxxxx, respectively. Supplementary data, including full annotation results, Augustus gene prediction mode, full assembly statistics, and repetitive elements library, can be accessed from the Open Science Framework (OSF) repository at https://doi.org/10.17605/OSF.IO/CE2AB.

